# Less is more: Light sampling via throttled visual phototransduction robustly synchronizes the *Drosophila* circadian clock in the absence of Cryptochrome

**DOI:** 10.1101/2020.03.10.985044

**Authors:** Maite Ogueta, Roger C Hardie, Ralf Stanewsky

**Author notes:** Corresponding author and Lead Contact, Ralf Stanewsky.

## Abstract

The daily changes of light and dark exemplify a prominent cue for the synchronization of internal circadian clocks to external time. The match between external and internal time is crucial for the fitness of organisms and desynchronization has been linked to numerous physical and mental health problems in humans. Organisms therefore developed complex and not fully understood mechanisms to synchronize their circadian clock to light. In mammals and in *Drosophila* both the visual system and dedicated non-image forming photoreceptors contribute to light resetting of the circadian clock. In the fruit fly, light-dependent degradation of the clock protein TIMELESS (TIM) by the blue light photoreceptor Cryptochrome is considered the main mechanism for clock synchronization, although the visual system also contributes. In order to understand the nature of the visual system contribution, we generated a genetic variant exhibiting extremely slow phototransduction kinetics, yet normal sensitivity. We show that in this variant the visual system is able to contribute its full share to circadian clock entrainment, both with regard to behavioral and molecular synchronization to light:dark cycles. This function depends on an alternative Phospholipase C-ß enzyme, encoded by *PLC21C*, presumably playing a dedicated role in clock resetting by light. We show that this pathway requires the ubiquitin ligase CULLIN-3, presumably mediating CRY-independent degradation of TIM during light:dark cycles. Our results suggest that visual system contribution to circadian clock entrainment operates on a drastically slower time scale compared with fast, visual and image forming phototransduction. Our findings are therefore consistent with the general idea that the visual system samples light over prolonged periods of time (hours) in order to reliably synchronize their internal clocks with the external time.

## Introduction

Physiology and behavior are temporally organized on seasonal and daily time scales by the interplay between the external environment and internal endogenous clocks. Circadian clocks influence both daily and seasonal rhythms and are synchronized with the external time through daily changes of light and darkness and the accompanying temperature cycles [1]. Desynchronization between external and internal time reduces overall fitness and is linked to several physiological and mental disorders in humans [2–4]. It is therefore important to understand the mechanisms underlying clock synchronization with the natural environmental rhythms. Both, in mammals and in the fruit fly *Drosophila melanogaster*, the visual image forming system as well as dedicated photoreceptor cells and photopigments contribute to light synchronization of the circadian clock [5–7]. In the fruit fly, synchronization to light:dark (LD) cycles is mediated by the visual system and by Cryptochrome (CRY), a blue light photoreceptor expressed within most clock cells [5,7–11]. Interestingly, the visual system can synchronize the clock using two different Rhodopsin signaling cascades: The canonical pathway, in which light-activated Rhodopsin activates Gq, resulting in activation of Phospholipase C-ß (PLC-ß), encoded by the *norpA* gene, and an alternative pathway, activating a PLC-ß encoded by the *Plc21C* gene [7]. While the canonical, *norpA*-dependent pathway synchronizes the clock at low (5-10 lux) light-intensities involving Rh1, Rh3, Rh4 and Rh6 signaling [9], the *Plc21C*-dependent pathway operates at higher intensities (~180 lux) and in complete absence of *norpA* function, involving at least Rh1, Rh5, and Rh6 [7].

Molecularly, the fly clock is reset by the light-dependent degradation of the clock protein Timeless (TIM). Together with Period (PER), TIM functions as a repressor of *per* and *tim* transcription, by inhibiting the positive transcription factors Clock (CLK) and Cycle (CYC) [12]. Only after degradation of PER and TIM, can CLK and CYC initiate another round of *per* and *tim* transcription, resulting in 24-hr molecular rhythms, which ultimately drive behavioral rhythmicity. Because PER is stabilized by binding to TIM, light-dependent TIM degradation also results in low PER levels, thereby removing both repressor proteins and enabling CLK and CYC to activate *per* and *tim* transcription. This constitutes an elegant way to synchronize the molecular oscillations with the external light conditions: Light in the early evening will delay the clock, because PER and TIM degradation can be compensated by translation of *per* and *tim* mRNA, which is at peak levels at this time of day. In contrast, light late at night will advance the clock, because PER and TIM levels remain low after degradation, due to trough levels of *per* and *tim* mRNA in the early morning.

Locomotor activity rhythms in *Drosophila* are controlled by approximately 150 clock neurons in the fly brain, characterized by rhythmic clock gene expression described above [13]. Interestingly about 50 % of the clock neurons express CRY and are therefore directly photosensitive [14]. Upon light exposure CRY undergoes a conformational change allowing it to bind to TIM, which triggers degradation of TIM, CRY and PER in the proteasome [15–17]. Nevertheless, CRY-independent mechanisms for light-dependent TIM degradation must exist, because molecular and behavioral resetting occurs in the absence of CRY [7,18]. In fact, TIM is degraded in artificially activated clock neurons in a CRY-independent manner [19], suggesting that the same Cullin-3 dependent mechanism could operate in clock neurons activated by light from the visual system. Moreover, it was recently shown that most of the clock neurons depolarize in response to brief (few ms) light pulses and that these responses depend on the visual system and not on Cry [20]. While the anatomical connections between the visual system and the circadian clock neurons remain elusive, it is clear that retinal photoreception plays an important role in molecular and behavioral resetting of the circadian clock to light.

Interestingly, the *norpA*-dependent and *norpA*-independent pathways contributing to light synchronization of the clock, also contribute to ERG responses of retinal photoreceptor cells to brief flashes of light [7]. Previously, it was shown that the almost complete lack of ERG responses in *norpA* mutants could be partially restored by simultaneous mutation of the *rdgA* gene, encoding DAG-kinase [21,22]. Due to the similar effects on ERG responses and circadian synchronisation to light, we investigated if removal of *rdgA* function would also improve clock restting in viusally impaired and CRY-deficient *norpA*^*P41*^ *cry*^*02*^ mutants. Surprisingly, we indeed observed a dramatic rescue of visual contribution to clock resetting and ERG sensitivity by simultanous removal of *norpA* and *rdgA* function. This remarkable function of a ‘throttled’ visual system depends on Rhodopsin phototransduction via *Plc21C*. While the kinetics of ERG responses in *norpA* and *rdgA* double mutants is orders of magnitude slower, behavioral clock-resetting speed and molecular synchronizaton of PER oscillations are basically identical to that of *cry* mutants, suggesting that circadian light-resetting depends on light-sampling over long time intervals. Finally we present evidence that visual sysem input requires CUL-3 function, suggesting that both CUL-3 and CRY-dependent TIM degradation contribute to light-synchronization of the *Drosophila* clock.

## Results

### *norpA rdgA* double mutants display normal light sensitivity combined with slow phototransduction

Previously, we found that tiny residual light responses can still be detected in photoreceptor whole-cell recordings from severe, and even supposedly null *norpA* alleles, and that these can be massively facilitated by mutations in the *rdgA* gene encoding DAG kinase [21,22]. For example although only minimal (~1-5pA) responses could be elicited by bright flashes in a supposedly null allele (*norpA*^*P24*^), responses to light in *norpA*^*P24*^ *rdgA*^*1*^ can reach several 100 pA [21]. The origin of the residual responses in *norpA*^*P24*^ mutants remained unresolved, and it could not be excluded that the *norpA*^*P24*^ mutation (which potentially leaves a truncated protein with intact catalytic site) is not truly null.

Here we explored sensitivity *in vivo* in the null *norpA*^*P41*^ allele [11] using electroretinogram (ERG) recordings from intact flies. As previously reported [7] responses in *norpA*^*P41*^ mutants were very small with < 1mV responses to the brightest test intensities (equivalent to ~ 10^7^ photoisomerisations per 1 sec flash) reflecting the requirement for *norpA* PLC for normal phototransduction [23]. Responses in *rdgA*^*1*^ mutants were also very severely reduced (Figure 1 and see [24]), this time reflecting the severe early onset and light independent retinal degeneration in this mutant (Figure S1). This degeneration is believed to be due to the cytotoxic effects of Ca^2+^ influx via constitutive activation of the light-sensitive TRP and TRPL channels during pupal development which prevents development of the microvillar rhabdomeres [24].

**Figure 1.**
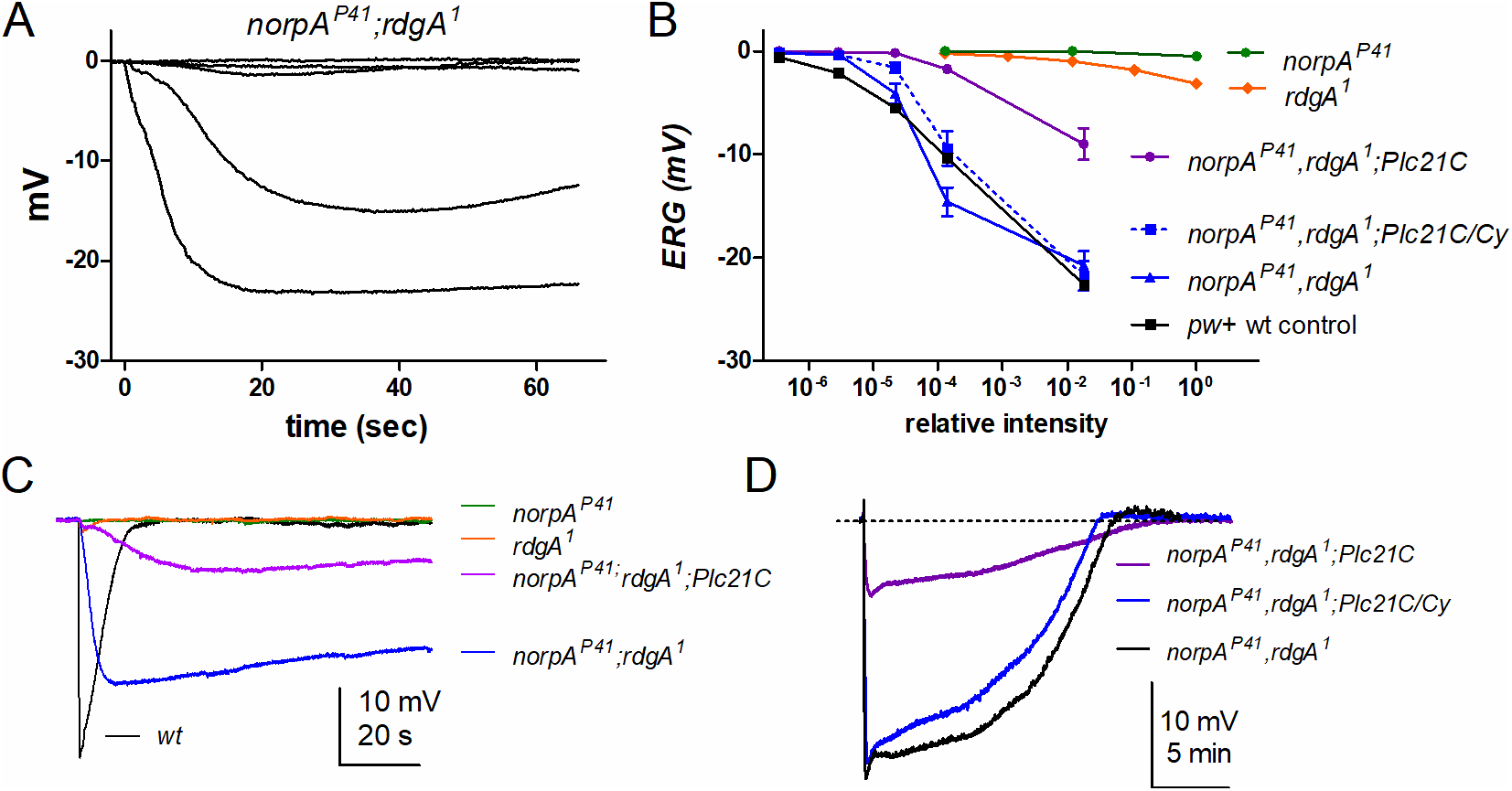
*norpA rdgA* double mutants display normal light sensitivity in their ERG response to a light flash. A) Responses in dark-reared *norpA*^*P41*^,*rdgA*^*1*^;*cry*^*02*^ flies to flashes (1s) of increasing intensity (relative intensities: 10^−6^ - 10^−2^). B) Summary of response intensity function data: single *norpA*^*P41*^ and *rdgA*^*1*^ mutants, all other genotypes were in *cry^02^* background. *norpA*^*P41*^*rdgA*^*1*^;*Plc21C^P319^*/+ were additional controls (same-aged siblings) for *norpA*^*P41*^*rdgA*^*1*^;*Plc21C*^*P319*^ flies. Mean ± SEM n = 8-12 flies for each genotype. C) Examples of responses in *norpA*^*P41*^ and *rdgA*^*1*^ single mutants, *norpA*^*P41*^ *rdgA*^*1*^ double mutants, and *norpA*^*P41*^ *rdgA*^*1*^;*Plc21C*^*P319*^ treble mutants to saturating 1 second flashes. D) Responses in *norpA*^*P41*^ *rdgA*^*1*^;*Plc21C*^*P319*^ compared to *norpA*^*P41*^ *rdgA*^*1*^ and *norpA*^*P41*^ *rdgA*^*1*^;*Plc21C^P319^/+* sibling controls on longer timescale.

The severe, early retinal degeneration in *rdgA*^*1*^ is completely suppressed in the *norpA*^*P24*^ *rdgA*^*1*^ double mutant, presumably because *norpA* PLC is required for the constitutive activation of the channels[21,24,25]. Correspondingly, we found a complete rescue of retinal degeneration in dark-reared *norpA*^*P41*^ *rdgA*^*1*^ double mutants with fully intact rhabdomeres as viewed in the deep pseudopupil or by optical neutralization, and near wild-type rhodopsin levels as determined by spectrophotometry (Figure S1).

To test sensitivity to light, we recorded the ERG in response to 1 second flashes of increasing intensity. Remarkably, the resulting response intensity functions from *norpA*^*P41*^ *rdgA*^*1*^ double mutants were now virtually indistinguishable from wild-type controls over a 5 log unit range, in both cases reaching amplitudes in excess of 20 mV (Figure 1B). Nevertheless, responses from *norpA*^*P41*^ *rdgA*^*1*^ showed profound differences in kinetics. Whilst responses in wild-type reach peak values within ~ 500 ms and return to baseline within 5-10 seconds, responses in *norpA*^*P41*^ *rdgA*^*1*^ peaked only after ~20-50s and then returned very slowly to baseline over many minutes (Figure 1C, D). As previously discussed, the slow rise and decay of the response in *norpA* backgrounds is believed to reflect the slow rate of encounters between activated Gq alpha subunits (which remain active indefinitely) and, the now, very rare residual PLC molecules [22,26,27]. The slow kinetics also necessitates leaving flies for 10-20 minutes in the dark between each test flash in order to reveal the true sensitivity to light.

Because *norpA*^*P41*^ is believed to be a genuine null allele [11], we asked whether light responses in *norpA*^*P41*^ *rdgA*^*1*^ might derive from an alternative PLC isoform. The *Drosophila* genome contains only one other PLC beta isoform, namely Plc21C. We therefore recorded ERGs in a triple mutant *norpA*^*P41*^ *rdgA*^*1*^ Plc21C using the hypomorphic *Plc21C*^*P319*^ allele [28]. Although sizeable (~10 mV) responses could still be recorded in this treble mutant, sensitivity was very severely (~ 100-fold) attenuated with respect to *norpA*^*P41*^ *rdgA*^*1*^ and *norpA*^*P41*^ *rdgA*^*1*^ *Plc21C*/+ sibling controls (Figure 1 B-D). This strongly suggests that Plc21C is responsible for the large, slow, responses in the *norpA*^*P41*^ *rdgA*^*1*^ double mutant.

### Lack of *rdgA* function rescues behavioral synchronization of *norpA*^*P41*^ *cry*^*02*^ double mutants to light:dark cycles

We previously showed that the results obtained from ERG responses to brief light exposure correlate with behavioral synchronization to ramping 12 hr : 12 hr light-dark (LD) cycles [7]. We therefore investigated if the dramatic rescue of *norpA*^*P41*^ ERG light-responses in *norpA*^*P41*^ *rdgA*^*1*^ double mutants also extends to circadian clock synchronization during ramping LD cycles (180 lux). First we analyzed the behavior of wild type control flies, as well as that of single mutant *norpA*^*P41*^, *rdgA*^*1*^, *cry^02^*, double mutant *norpA*^*P41*^ *rdgA*^*1*^, *norpA*^*P41*^ *cry^02^*, *rdgA*^*1*^ *cry^02^* as well as treble mutant *norpA*^*P41*^ *rdgA*^*1*^ *cry^02^* flies in LD cycles, followed by exposure to constant darkness (Figure 2 and S2A). Control flies exhibited typical bimodal behavior with two prominent activity peaks in the morning and evening, whereby the activity starts to increase several hours before the respective light transitions (Figure 2A, B). This behavioral anticipation of changes in the environmental condition is controlled by the circadian clock and indicates proper synchronization to LD [29]. Similar to the control flies, all single mutants and *norpA*^*P41*^ *rdgA*^*1*^ double mutants synchronized well to LD cycles (Figure 2, S2A), confirming that both light input via visual system and Cry contribute to synchronization of the circadian clock [5,10]. Similarly *norpA*^*P41*^ *cry^02^* double mutants showed almost normal bimodal behavior (Figure S2A), confirming that *norpA*-independent visual photoreception also contributes to circadian clock synchronization [7,11]. In contrast, double mutant *rdgA*^*1*^ *cry^02^* flies exhibited a less pronounced bimodal activity pattern with high activity levels during the day and night (Figure 2A, B). This indicates that light synchronization in *rdgA*^*1*^ *cry^02^* double mutants is impaired, presumably because the retinal degeneration caused by *rdgA*^*1*^ affects both *norpA*-dependent and *norpA*-independent photoreception. Strikingly, the behavior of *norpA*^*P41*^ *rdgA*^*1*^ *cry^02^* treble mutants was indistinguishable from that of wild type controls, indicating a rescue of retinal degeneration and perhaps even an improvement compared to *norpA*^*P41*^ *cry^02^* double mutants (Figure 2A, B). As expected, flies from all genotypes showed robust behavioral rhythmicity in the final DD part of the experiments, indicating that none of the affected genes is required for circadian clock function (Figure 2A, Table S1).

**Figure 2.**
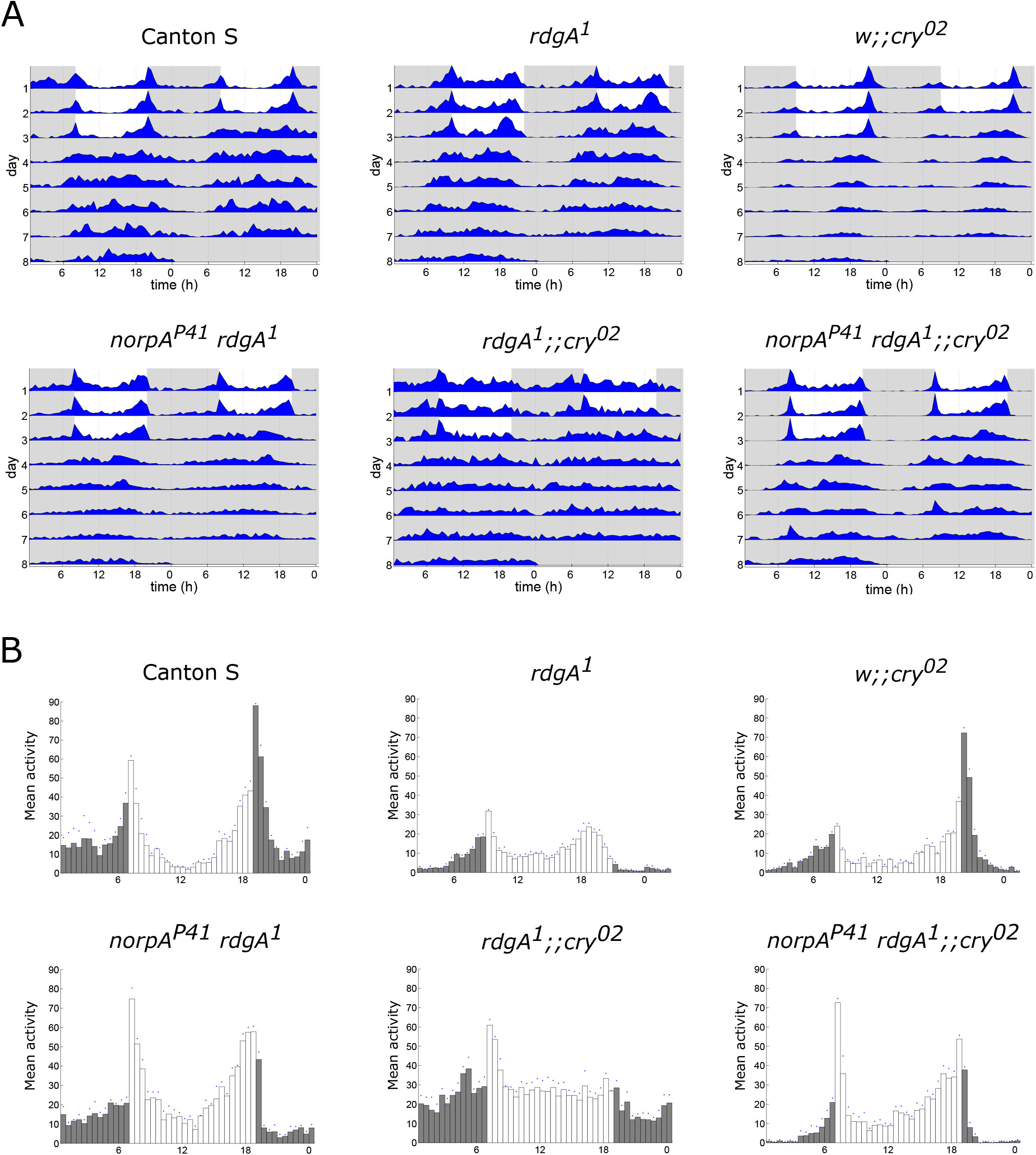
Loss of *norpA* function restores normal LD behavior in *rdgA*^*1*^ *cry^02^* double mutants. Average actograms (A) and histograms (B) of each of the indicated genotypes (n = 16). Flies were exposed to LD for 3 days, followed by 5 days of constant darkness (DD) at 25°C.White areas indicate lights-on, grey areas lights-off. In (B) the average activity of the 3 days in LD is shown.

To directly test the effects of the various mutant combinations on light synchronization we performed ‘jet-lag’ experiments, in which flies were first exposed to a combined LD and temperature cycle (TC), followed by an LD cycle that was phase delayed by 6 hours compared to the initial LDTC. The TC was in phase with the LD cycle, to ensure that genotypes with impaired light inputs would nevertheless be synchronized before being exposed to the phase delayed LD cycle [7]. As expected from the LD experiments described above, control flies and single mutants synchronized to the initial LDTC and resynchronized quickly to the phased-delayed LD cycle (Figure 3). Quantification of the number of days required for resynchronization revealed that *cry^02^* flies need 3 days to adjust to the phase-delayed LD cycle, while control and *norpA*^*P41*^ and *rdgA*^*1*^ single and double mutants need only 2-3 days (Figure 3B; S2B; Table 1;[7]). Interestingly, *rdgA*^*1*^ single mutants exhibit an earlier evening phase during LDTC and LD cycles, which is maintained during constant conditions (Figures 2; 3A, B). To test, if the earlier evening phase is indeed caused by the impaired *rdgA* function, we tested the strong hypomorphic allele *rdgA^KS60^* [30], which indeed showed the same advance of the evening peak, also observed in transheterozygous *rdgA^1^/rdgA^KS60^* females (Figure S2A, B). Because both *rdgA* alleles do not affect free-running period length (Table S1), and *rdgA*^*1*^ resynchronizes normally to shifted LD cycles (Figure 3), *rdgA* is required for a normal phase of the evening activity peak during LD. Interestingly, the phase advance is rescued in the *norpA*^*P41*^ *rdgA*^*1*^ double mutants (Figure S2B), indicating that it is related to retinal degeneration observed in the *rdgA* single mutants (Figure S1). As previously reported, *norpA*^*P41*^ *cry^02^* double mutants require 5 days for resynchronization (Figure 3A, C, Table 1) [7]. Surprisingly, and consistent with the results from the LD experiment, *rdgA*^*1*^ *cry^02^* double mutants completely fail to synchronize to the phase-delayed LD cycle (Figure 3A, C; Table 1). Strikingly, light synchronization is completely restored in *norpA*^*P41*^ *rdgA*^*1*^ *cry^02^* treble mutants, which require only 3 days for resynchronization, similar to *cry^02^* single mutants (Figure 3A-C, Table 1). Analysis of the activity peak phase at day 1 and day 2 after the shift of the light dark cycle revealed that resynchronization of *norpA*^*P41*^ *rdgA*^*1*^ *cry^02^* flies is significantly faster compared to *norpA*^*P41*^ *cry^02^* double mutants. This confirms that removal of *rdgA* function somehow rescues the lack of *norpA* function in light-signaling to the clock and vice versa (Figure 4A, B). Moreover, on day 2 after the light shift the phase of *norpA*^*P41*^ *rdgA*^*1*^ *cry^02^* flies is indistinguishable from that of single mutant *cry^02^* flies, indicating that parallel removal of *norpA* and *rdgA* function almost completely restores visual system input to the circadian clock (Figure 4B).

**Figure 3.**
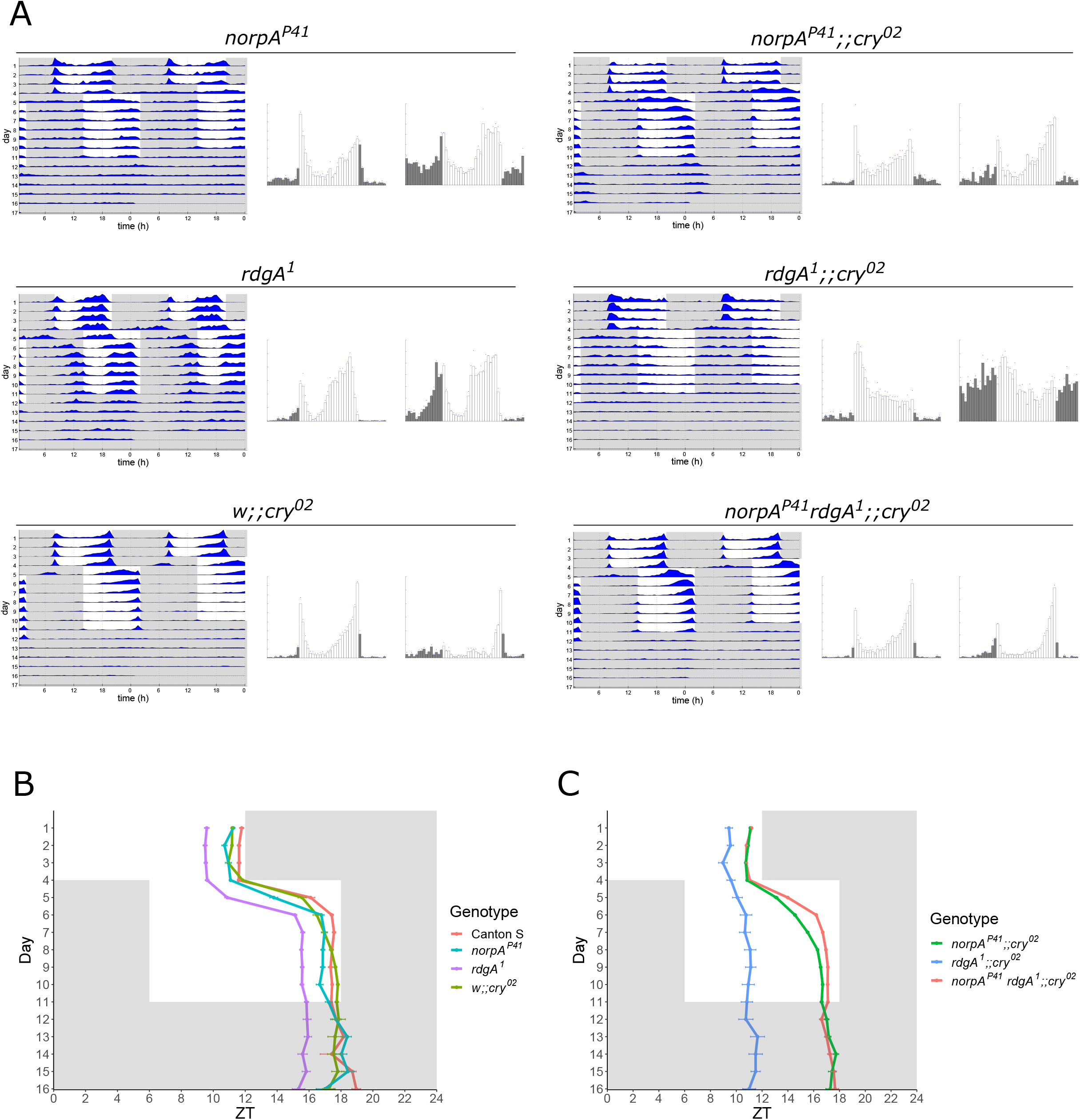
Reciprocal rescue of CRY-independent light resetting in *norpA*^*P41*^ and *rdgA*^*1*^ double mutants. Flies were entrained to a 2-hr ramping LD cycle in combination with temperature cycles of 25:16°C. After 4 days, the temperature was kept constant at 25°C, and the LD regime was delayed by 6 hr (‘jetlag’). The flies were kept in these new conditions for 7 days, followed by 5 days of DD. (A) Average actograms (left) depicting activity throughout the experiment, and histograms averaging the activity during the 3 days before the shift (middle) and the last 3 days after the shift (right) (n = 16 for each genotype). Shading as in Figure 2. (B-C) Quantification of the evening activity peak phase for each day of the experiment for single mutants (B), or mutant allele combinations (C). See also Table 1.

**Table 1.** Number of days required for resynchronization to a phase-delayed LD cycle. The number of days the different genotypes need to re-entrain was calculated by counting the days until each fly reached a stable phase relationship with the new LD cycle. The ‘Δ hours’ result from subtracting the phase at the last day of the first LD (day 4) from that of the last day after the shift (day 11).

| Genotype | Day to re-entrain | Δ Hours | n |
| --- | --- | --- | --- |
| Canton S | 1.5 ± 0.1 | 5.8 ± 0.1 | 32 |
| *norpA<sup>P41</sup>* | 2.4 ± 0.1 | 6.1 ± 0.2 | 32 |
| *w;;cry<sup>02</sup>* | 3.1 ± 0.2 | 5.9 ± 0.1 | 32 |
| *norpA<sup>P41</sup>;;cry<sup>02</sup>* | 5.7 ± 0.5 | 5.8 ± 0.1 | 158 |
| *rdgA<sup>1</sup>* | 2.4 ± 0.1 | 6.2 ± 0.1 | 64 |
| *rdgA<sup>1</sup>;;cry<sup>02</sup>* | - | 1.2 ± 0.4 | 54 |
| *norpA<sup>P41</sup> rdgA<sup>1</sup>* | 2.3 ± 0.1 | 6.1 ± 0.1 | 94 |
| *norpA<sup>P41</sup> rdgA<sup>1</sup>;;cry<sup>02</sup>* | 2.9 ± 0.1 | 6.1 ± 0.1 | 138 |
| *norpA<sup>P41</sup> rdgA<sup>1</sup>;PLC21C<sup>P319</sup>;cry<sup>02</sup>* | - | 2.5 ± 0.7 | 55 |
| *norpA<sup>P41</sup> rdgA<sup>1</sup>;Gq<sup>V303D</sup>;cry<sup>02</sup>* | - | 2.3 | 23 |
| *norpA<sup>P41</sup>;Clk856-Gal4; cry<sup>b</sup>* | 5.4 ± 0.2 | 5.6 ± 0.1 | 32 |
| *norpA<sup>P41</sup>;Clk856-Gal4;UAS-Cul3 RNAi cry<sup>b</sup>* | 6.4 ± 0.2 | 5.8 ± 0.3 | 16 |
| *norpA<sup>P41</sup>;elav-Gal4;UAS-Cul3 RNAi cry<sup>b</sup>/cry<sup>02</sup>* | - | 0.6 ± 0.2 | 13 |

**Figure 4.**
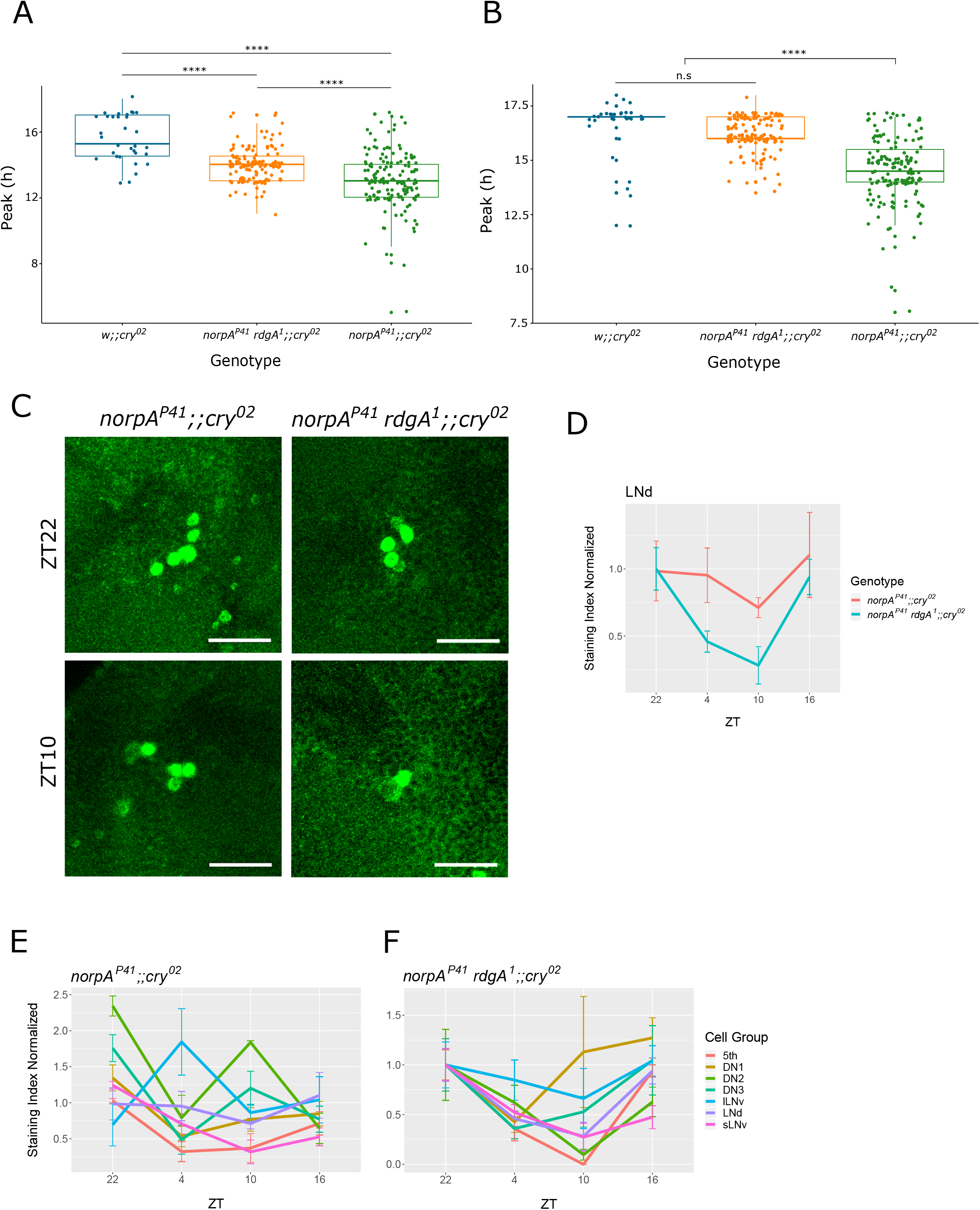
Significant improvement of behavioral and molecular resynchronization in *norpA*^*P41*^ *cry^02^* double mutants by simultaneous removal of *rdgA* function. Boxplots representing the position of the activity peak on day one (A) and day two (B) after the shift. Middle line represents the median, bottom and top borders the first and third quartile respectively, and whiskers indicate minimum and maximum values. The single dots correspond to the values of single flies. (A) On day one after the shift, the peak of *cry^02^* is already at 15.5 ± 0.2 hr, while that of *norpA*^*P41*^ *cry^02^* is at 13.1 ± 0.1 hr and that of *norpA*^*P41*^ *rdgA*^*1*^ *cry^02^* is in between at 14.0 ± 0.1 hr. All groups are statistically significant from another with p<0.001. (B) Two days after the shift, there is no significant difference between *wcry^02^* and *norpA*^*P41*^ *rdgA*^*1*^ *cry^02^* (16.5 ± 0.2 vs 16.2 ± 0.1 hr respectively), while in *norpA*^*P41*^ *cry^02^* the peak of the activity occurs significantly earlier (14.5 ± 0.1 hr) (p<0.001). (C) anti-PER labelling of the LNd on day 2 of the phase-delayed LD cycle. Consistently, only 1 PER^+^ LNd was detected at ZT10 in *norpA*^*P41*^ *rdgA*^*1*^ *cry^02^* brains, indicating that most LNd are synchronized by the visual system in this group. Scale bar: 20μm. (D) Quantification of PER expression in LNd at four different time points. (E, F) quantification of all other clock neuronal groups in *norpA*^*P41*^ *cry^02^* (E), and *norpA*^*P41*^ *rdgA*^*1*^ *cry^02^* flies.

### Lack of *rdgA* function rescues molecular synchronization of PER expression in clock neurons of *norpA*^*P41*^ *cry*^*02*^ double mutants

To determine if removal of *rdgA* function also improves molecular synchronization of *norpA*^*P41*^ *cry^02^* double mutants we analyzed PER expression in clock neurons during the same phase-delayed LD cycle. Brains of *norpA*^*P41*^ *cry^02^* and *rdgA*^*1*^ *norpA*^*P41*^ *cry^02^* mutants were dissected at 4 time points during the 2^nd^ day after the light shift allowing direct comparison to the behavioral phase-determination performed on the same day (Figure 4 B). As expected from the behavioral results, *rdgA*^*1*^ *norpA*^*P41*^ *cry^02^* treble mutants displayed an overall improvement of PER synchronization in the clock neurons (Figure 4 C-F). In particular, in the LNd, where PER expression is completely desynchronized in the *norpA*^*P41*^ *cry^02^* double mutants, PER cycling is restored in the treble mutant with expected peaks and troughs at ZT22 and ZT4, respectively (Figure 4 C-F). In addition, in *rdgA*^*1*^ *norpA*^*P41*^ *cry^02^* brains most other neuronal groups showed restored PER cycling with trough expression at ZT10 and peak expression between ZT16 and ZT22 (Figure 4 E,F; S3). Exceptions are the CRY-dependent l-LNv [10], as well as the DN1 and DN3, where trough expression remains at ZT4 as in the *norpA*^*P41*^ *cry^02^* double mutants. Also, biphasic PER oscillations observed in the DN2 and DN3 of the double mutants, indicating desynchronization between neurons, were not observed in treble mutants (Figure 4 E,F, S3), further indicating improved molecular synchronization to LD cycles in the absence of *rdgA* function.

### Loss of *norpA* and *rdgA* function does not compensate for the lack of Cryptochrome

Although the behavior during phase-delayed LD cycles indicated that *norpA*^*P41*^ *rdgA*^*1*^ *cry^02^* treble mutants synchronize to light no faster than *cry^02^* single mutants do (Figure 3), we wanted to test the possibility that throttled phototransduction due to the loss of *norpA* and *rdgA* function may compensate the lack of CRY. For this, we exposed control flies, the *norpA*^*P41*^ *rdgA*^*1*^ *cry^02^* treble mutants, and *cry^02^* flies to 30 min light pulses (LP) at ZT15 and ZT22 on the last day of an LD cycle. As expected, a ZT15 LP resulted in ~ 3 hr phase delays of the activity onset during the constant conditions following LD (Figure S4), while a ZT22 LP induced ~1.5 hr phase advances (Figure S4) [10]. Also as expected, single *cry^02^* mutants did not display any significant phase delays or advances, and the same was true for the *norpA*^*P41*^ *rdgA*^*1*^ *cry^02^* treble mutants (Figure S4).

### *PLC21C* is responsible for circadian clock synchronization in *norpA^P41^ rdgA^1^ cry^02^* flies

The robust ERG responses observed in *norpA*^*P41*^ *rdgA*^*1*^ double mutants depend on the PLC-ß encoded by *Plc21C* (Figure 1 B-D). Moreover, we have shown that *Gq-Plc21C* signaling mediates *norpA*-independent light-input to synchronize the circadian clock [7]. We therefore tested if *Plc21C* is also required for the robust synchronization of *norpA*^*P41*^ *rdgA*^*1*^ *cry^02^* mutants to LD cycles. For this, we generated *norpA*^*P41*^ *rdgA*^*1*^ *Plc21C*^*P319*^ *cry^02^* mutants and analyzed their behavior in plain LD cycles and in jetlag conditions (Figure 5). The quadruple mutants did show an activity peak during the 2^nd^ half of the warm phase during LDTC, indicating that they can synchronize to combined LD and temperature cycles (Figure 5A). After the 6 hour delay of the LD cycle during constant temperature, locomotor activity spread out more evenly between day and night, lacking any distinct peaks or troughs (Figure 5A). The flies did clearly not resynchronizing to the delayed LD cycle and became largely arrhythmic in constant conditions following exposure to LD (Figure 5A,C; Table 1, S1).

**Figure 5.**
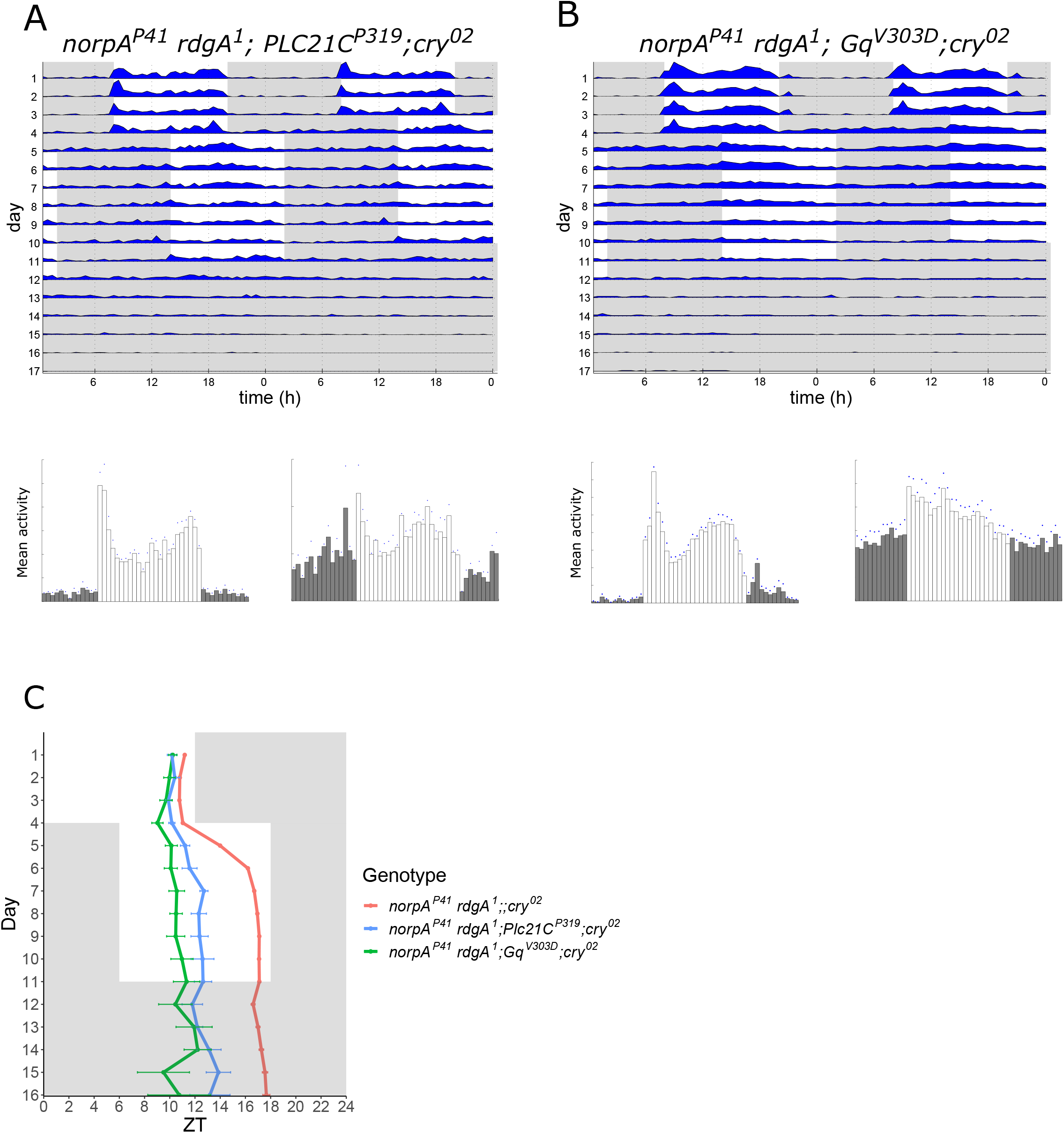
PLC21C and Gq mediate visual system resetting in the absence of *norpA* and *rdgA* function. (A, B) Representative average actograms (top) of 16 *norpA*^*P41*^ *rdgA*^*1*^;*Plc21C*^*P319*^;*cry^02^* (A) and 17 *norpA*^*P41*^ *rdgA*^*1*^;*Gq^V303D^*;*cry^02^* flies during a ‘jetlag’ experiment (for detailed conditions see Figure 3). The histograms below represent average activity of the same flies 3 days before the shift (left) and the 3 last days in LD after the shift (right). (C) Quantification of the position of the evening peak in *norpA*^*P41*^ *rdgA*^*1*^;;*cry^02^* (n=138), *norpA*^*P41*^ *rdgA*^*1*^;*Plc21C*^*P319*^;*cry^02^* (n=55), and *norpA*^*P41*^ *rdgA*^*1*^;*Gq*^*V303D*^;*cry^02^* (n=23) flies.

To analyze the involvement of *Gq*, we generated *norpA*^*P41*^ *rdgA*^*1*^ *Gq*^*V303D*^ *cry^02^* mutants and analyzed their behavior under the same conditions. The flies behaved very similar to the *norpA*^*P41*^ *rdgA*^*1*^ *Plc21C*^*P319*^ *cry^02^* mutants, showing synchronization to the initial LDTC cycle, but were not able to adjust their activity to the phase-delayed LD cycle (Figure 5B,C, Table 1). Moreover, they also exhibited increased arrhythmicity in constant conditions (Figure 5B, Table S1). The results show that visual system function in the absence of *norpA* and *rdgA* requires *Gq* and *PLC21C* for robust clock synchronization.

### The ubiquitin ligase CULLIN-3 is required for light-resetting in *norpA*^*P41*^ *cry*^*b*^ mutants

To identify the molecular mechanism of clock synchronization in the absence of CRY signaling, we investigated the potential involvement of CULLIN-3 (CUL-3), because TIM is a known target of this ubiquitin ligase after neuronal activation and in constant conditions [19,31]. For this we combined an established *Cul-3* RNAi line [31] with *cry*^*b*^ by meiotic recombination and subsequently generated a *norpA*^*P41*^ *Cul-3* RNAi *cry*^*b*^ stock. These flies were crossed to *norpA*^*P41*^ *cry*^*b*^ and *norpA*^*P41*^ *cry^02^* flies, carrying the pan-neuronal driver lines *elav-gal4* or the clock neuronal driver *Clk856-gal4*, respectively. The resulting progeny, mutant for *norpA* and *cry*, and expressing *Cul-3* RNAi in all neurons, or all clock neurons, where then tested for resynchronization to LD cycles using the jetlag assay described above. In contrast to the controls, *norpA*^*P41*^ *cry*^*b*^ flies with pan-neuronal RNAi-mediated CUL-3 knockdown failed to re-synchronize to the phase-delayed LD cycle and maintained an evening peak phase established during the initial LDTC regime (Figure 6A, C, Table 1). Similarly, CUL-3 knockdown in all clock neurons resulted in slower re-synchronization compared to *norpA*^*P41*^ *cry^02^* control flies (Figure 6A, B, Table 1). The weaker phenotype after *Clk856*- compared to *elav*-mediated CUL-3 knock down, could be due to differences in driver strength in combination with the efficiency of the RNAi-construct. Nevertheless, the results demonstrate a prominent role for CUL-3 in light-resetting, supporting the idea that this ubiquitin ligase mediates light-dependent TIM degradation in the absence of CRY.

**Figure 6.**
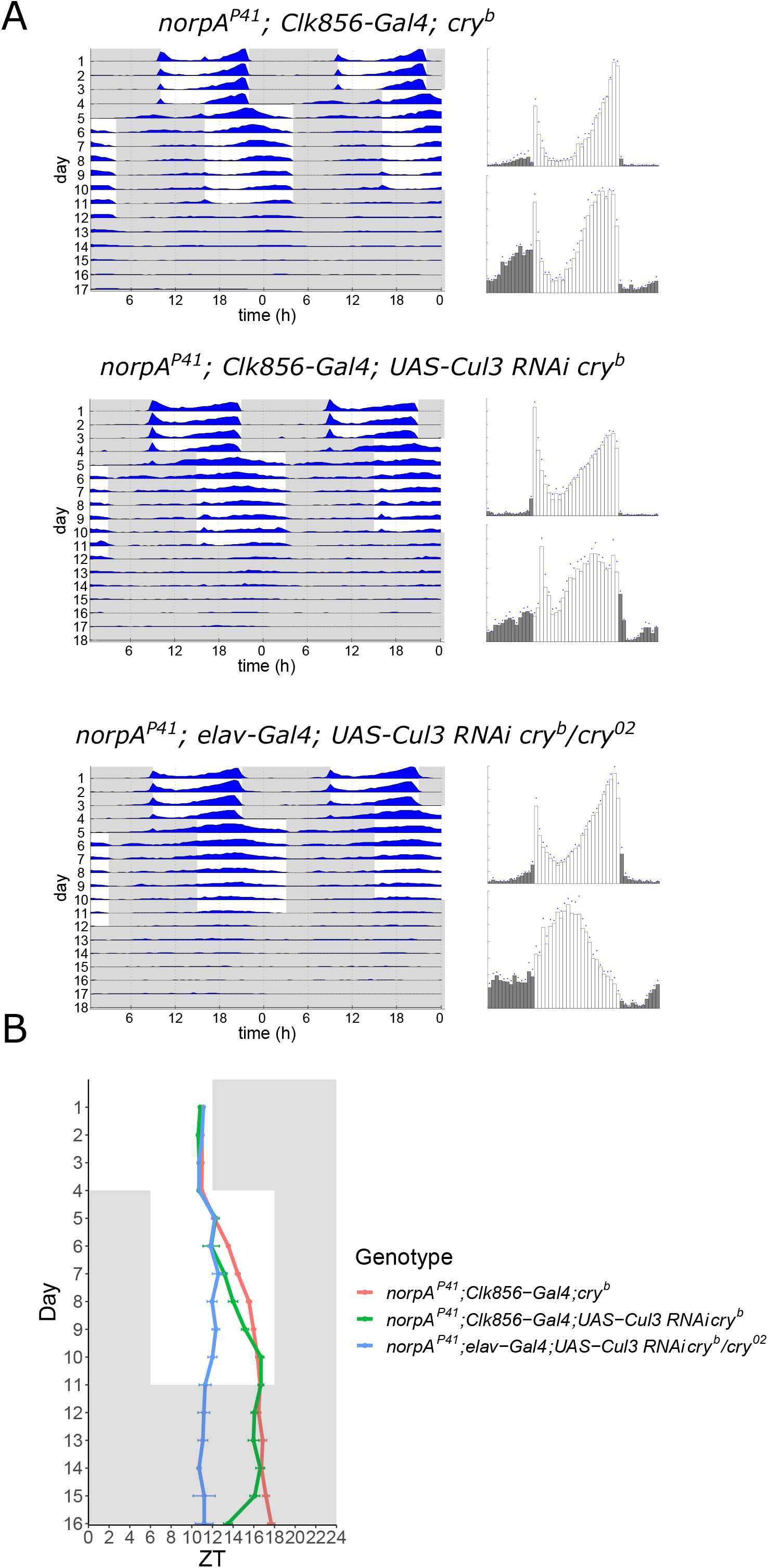
*Cullin-3* is mediates *cry*- and *norpA*-independent synchronization to light:dark cycles. (A) Average actograms (left) and histograms (right) of the indicated actograms in a “jetlag” experiment using the same conditions as in Figure 3. (B) Quantification of the evening activity peak phase of the different genotypes. See also Table 1.

### *rdgA*^*1*^ encodes a glycine to serine change in the catalytic domain of DAG-kinase

The *rdgA*^*1*^ allele was originally reported as an EMS-induced C -> T nucleotide change resulting in the insertion of a STOP codon at residue 1153, between the catalytic domain and ankyrin repeats of DAG-kinase [32]. While generating the *norpA*^*P41*^ *rdgA*^*1*^ double mutants, we noticed that our *rdgA*^*1*^ stock did not carry the expected nucleotide change. In addition, it did not carry any of the published nucleotide changes associated with other *rdgA* alleles (data not shown). Because our *rdgA*^*1*^ stock showed the expected ERG and retinal degeneration phenotypes (Figures 1C, S1), we decided to sequence the open reading frame of this allele. We found a G to A nucleotide change at nucleotide position 2928 (according to[32]), resulting in a Ser to Gly substitution at residue 897, which is a highly conserved position in the catalytic domain of DAG-kinase (Figure S5). The mutation is therefore 28 residues downstream of the Gly to Asp substitution encoded by *rdgA^2^*, a loss-of-function allele, which also maps to the catalytic domain [32]. Because the mutation in our *rdgA*^*1*^ stock maps to the catalytic domain and does not correspond to any other known *rdgA* mutations, the originally published *rdgA*^*1*^ sequence is most likely incorrect [32].

## Discussion

The Cry-centric view of circadian clock light-resetting has recently been challenged by the observation that visual system input elicits electrical responses in several clock neuronal groups independent of CRY [20]. Indeed, since CRY was identified as circadian photoreceptor in flies more than 20 years ago [10,33], it was clear that the visual system also plays an important role in clock resetting by light. Although CRY seems absolutely required for clock resetting to brief light pulses ([10,34,35], Figure S3), synchronization to more natural conditions of daily light:dark changes works well without CRY and depends on the presence of a functional visual system [5,10]. The results presented here emphasize the role of the visual system in circadian clock synchronization, and more importantly, shed light on the nature of its contribution.

### Slow phototransduction is sufficient for full visual system contribution to clock resetting

It was previously shown that in the complete absence of the visual system and CRY function the circadian clock can no longer be synchronized to light [5]. The observation that *rdgA*^*1*^ *cry^02^* double mutants do not synchronize to LD cycles is therefore not surprising, given the strong retinal degeneration phenotype induced by *rdgA*^*1*^ (Figure S1) [24]. It is also known that *norpA*^*P41*^ *cry*^*b*^ double mutants lead to severe decrements of light-resynchronization, more severe compared to the respective single mutants [10,36]. The slow resynchronization of *norpA*^*P41*^ *cry*^*b*^ double mutants could later be attributed to *norpA* independent Rhodopsin signaling involving a different PLC-ß enzyme, encoded by the gene *Plc21C* [7,11]. But how can the fast, *cry^02^*-like resynchronization kinetics of *norpA*^*P41*^ *rdgA*^*1*^ *cry^02^* treble mutants to LD cycles be explained? Although we observed a dramatic reciprocal rescue of the *rdgA*^*1*^ retinal degeneration and *norpA*^*P41*^ ERG phenotypes in *norpA*^*P41*^ *rdgA*^*1*^ flies, these double mutants lack crucial components of the canonical phototransduction cascade. *norpA*-encoded PLC-ß hydrolyses the membrane lipid PIP2 to the IP3 and DAG 2^nd^ messenger molecules, and the latter is one of several candidates implicated in direct and/or indirect activation of the TRP and TRPL channels [21,24,37–40]. *rdgA*-encoded DAG-kinase inactivates DAG by turning it into phosphatidic acid (PA), which is also the first step in the re-synthesis of PIP2 in the microvillar membrane [41]. Retinal degeneration in *rdgA*^*1*^ mutants is believed to be caused by constitutively open TRP and TRPL channels, possibly due to raised DAG and/or reduced PIP_2_ levels [42] and explains both the severe ERG phenotypes of *rdgA*^*1*^ single mutants and the impaired clock synchronization in *rdgA*^*1*^ *cry^02^* double mutants (Figures 1B,C; S1; 2, 3A,C). The simplest explanation for the retinal degeneration rescue of *rdgA*^*1*^ by *norpA*^*P41*^ would be the lower DAG levels, and/or restored PIP_2_ levels, because PIP_2_ hydrolysis is now limited to PLC21C activity. Conversely, the rescue of the *norpA*^*P41*^ light response by *rdgA*^*1*^ could in principle be explained on the basis that DAG is an excitatory 2^nd^ messenger, since the small amount of DAG generated by PLC21C would be expected to build up to higher levels in the absence of DAG kinase (for further discussion see [39]). Our results are consistent with the idea that based on the effects explained above the DAG levels in *norpA*^*P41*^ *rdgA*^*1*^ mutants are sufficient for normal sensitivity of the visual system photoreceptors to brief light pulses as well as for molecular and behavioral circadian synchronization to LD cycles. At the same time, they do not reach high enough levels to induce retinal degeneration.

The extremely slow ERG kinetics in response to brief (1 second) light pulses observed in the *norpA*^*P41*^ *rdgA*^*1*^ double mutants can probably be explained by the long times required for activated Gq alpha subunits to encounter and activate the very rare PLC21C molecules [22,26]. Importantly, this ‘slow’ or throttled phototransduction cascade seems perfectly suited for synchronizing the circadian clock to LD cycles, where light exposure occurs over many hours. This strongly supports the idea that for clock resetting under natural conditions the sampling of light over many hours plays an important role, both in insects and in the mammalian system [6,35]. It appears that in both systems the visual system has gained a light-sampling function independent from the fast image forming processes, which is dedicated to circadian clock synchronization. While in insects this may involve the recruitment of a specialized PLC-ß enzyme (PLC21C), in the mammalian system this involves specialized intrinsically photosensitive retinal ganglion cells (ipRGCs), expressing the invertebrate like melanopsin photopigment as well as rod and cone input into these cells [6,43]. With regard to invertebrate phototransduction, our results emphasize a role for DAG in TRP/TRPL channel opening.

### Role of PLC21C in the *Drosophila* clock

We confirm here the role of PLC21C for synchronization of the circadian clock to LD cycles ((Figure 5, [7]). Like in olfactory signal transduction, PLC21C light-resetting function requires Gq activation ([28], Figure 5) by Rhodopsin photopigments [7]. We show that in the absence of *norpA* and *rdgA* function, *Plc21C* mediates efficient Rhodopsin signaling to the clock neurons, so that resynchronization behavior is almost indistinguishable from that of flies with an intact visual system (Figures 3-5). In conjunction with the slow ERG kinetics of the photoreceptor cells after exposure to a light flash (Figure 1), this suggests that the normal function of PLC21C involves light-synchronization of the circadian clock, via sampling of light over long time intervals. In addition, PLC21C has been implicated to mediate behavioral responses to brief LP’s, presumably acting down stream of Rhodopsin 7 (Rh7) directly within the LNv clock neurons [44]. Because Rh7 null mutants rapidly synchronize to white-light LD cycles, and do not further slowdown resynchronization of *norpA*^*P41*^ *cry^02^* mutants [45], Rh7/PLC21C signaling appears to be both spatially (within LNv) and functionally (response to brief LP’s) distinct from Rhodopsin/PLC21C signaling in retinal photoreceptor cells. Interestingly, there is more precedence for PLC21C function within clock neurons, yet independent of light and downstream of G_o_ and not G_q_: Dahdahl et al [46] showed that PLC21C down regulation in LNv results in an 1-hr elongation of the free running period. Because LNv-expression of a constitutively active G_o_ protein reduced overall rhythmicity, the authors concluded that G_o_/PLC21C signaling in LNv is important for both rhythms strength and period length. This function may be related to the arrhythmicity we observed in the *norpA*^*P41*^ *rdgA*^*1*^ *Plc21C*^*P319*^ *cry^02^* mutants. But why did we not observe the expected longer period as would be expected from the results after PLC21C down-regulation [46]? We think that the poor synchronization of the quadruple mutant flies due to the presence of the visual mutations, may contribute to their arrhythmicity in constant conditions. In fact, the *norpA*^*P41*^ and *rdgA*^*1*^ single mutants combined with *cry^02^* synchronize poorly to LD cycles and show decreased rhythmicity in DD, while the *norpA*^*P41*^ *rdgA*^*1*^ *cry^02^* treble mutant synchronizes well to LD cycles and is strongly rhythmic in DD (Figures 3, 4; Table S1). Likewise mutants affecting synchronization to temperature cycles (TC), but not to LD cycles, showed decreased rhythmicity in constant conditions after TC and not after LD [47]. Our results therefore support a role for PLC21C downstream of Gq in retinal phototransduction and synchronization of the circadian clock to LD cycles, as well as downstream of G_o_ in clock function.

### Cryptochrome versus visual system mediated light synchronization of the circadian clock

Our findings highlight the importance of the visual system in clock synchronization and raise the question about the importance of CRY in this process. While it is clear that CRY is required for the response to brief light-pulses, this situation is presumably not relevant in natural conditions. For example, insects of the order hymenoptera (e.g., wasps, bees, and ants) and beetles lack the *Drosophila* type CRY and synchronize their clock presumably exclusively via the visual system [48]. The molecular mechanism of CRY-mediated clock synchronization is well understood (e.g. [15,17,49]), but we know little about the mechanism underlying molecular clock resetting in the absence of CRY. It has long been known that PER and TIM oscillations in subsets of the clock neurons can be synchronized by LD cycles in the absence of CRY, and our results implicate CUL-3 mediated TIM degradation as additional mechanism for molecular light resetting (Figure 6, Table 1). TIM is a known target for CUL-3 [31], and this ubiquitin ligase also mediates CRY-independent TIM degradation in artificially activated clock neurons [19]. Because it is known that light induces neuronal firing in most of the clock neurons and in the absence of CRY [20], our results, and in particular the improved molecular synchronization of most clock neurons in *norpA*^*P41*^ *rdgA*^*1*^ *cry^02^* mutants, suggest that the visual system resets the clock via neuronal activation, followed by CUL-3 mediated TIM degradation.

## Acknowledgements

We thank Gaiti Hasan and Francois Rouyer for fly stocks, and Luis Garcia for developing the programmable LED system. This work was supported by the European Union (H2020 Initial Training Network CINCHRON to R.S. and by the Marie Curie WHRI-Academy COFUND fellowship PCOFUND-GA-2013-608765 to M.O)

## Materials and Methods

### Flies

Flies were raised in a 12 h:12 h light dark (LD) cycle on standard Drosophila medium (0.7% agar, 1.0% soy flour, 8.0% polenta/maize, 1.8% yeast, 8.0% malt extract, 4.0% molasses, 0.8% propionic acid, 2.3% nipagen) at 25°C and 60% relative humidity. The hypomorphic *Plc21C*^*P319*^ and loss-of-function *norpA*^*P41*^ and *cry^02^* alleles have been described previously [11,28,50]. The *rdgA*^*1*^ and *rdgA^KS60^* alleles were described in [32]. The loss of function *Gq*^*V303D*^ allele is described in [51]. Here we combined *rdgA*^*1*^ with *norpA*^*P41*^ using meiotic recombination. Recombinants were confirmed by PCR (for *norpA*^*P41*^;[11]) and by the reappearance of the Deep Pseudopupil (DPP) (lacking in *rdgA*^*1*^, fully restored in the *norpA*^*P41*^ *rdgA*^*1*^ double mutants). Finally, *rdgA*^*1*^ was confirmed by sequencing, which lead to the correction of the published *rdgA*^*1*^ lesion (Figure S5, Table S2). The 3^rd^ chromosomal *UAS-Cul-3 RNAi* line was described in [31] and was combined with the *cry*^*b*^ *ss^1^* chromosome using meiotic recombination. Recombinants were isolated by screening for *w^+^* eye color (*UAS-Cul3-RNAi*) and *ss^1^* (maps close to *cry*) and confirmed by sequencing (*cry*^*b*^)

### Behavior

Analysis of locomotor activity of 4-5 day old male flies was performed using the Drosophila Activity Monitor System (DAM2, Trikinetics Inc., Waltham, MA, USA) with individual flies in recording tubes containing food (2% agar, 4% sucrose). For standard LD experiments, DAM2 activity monitors containing flies were located inside a light- and temperature-controlled incubator (Percival Scientific Inc., Perry, IA, USA), where fly activity was monitored for 4 days in rectangular 12hr:12h LD (400 lux generated by 17W F17T8/TL841 cool white Hg compact fluorescent lamps, Philips) followed by 5 days in constant darkness and temperature (25°C). For light resynchronization (jetlag) experiments, flies were also located inside a temperature controlled incubator, but light was provided by white LED strips controlled by an Arduino to control light intensity and to program lights on and off [52]. As described previously [7] lights were ramped for 2 hr at the beginning and end of each day. During the first 4 days, the LD cycles were combined with temperature cycles of 25°C during “lights-on” and 16°C during “lights-off” (LDTC), to ensure the entrainment of visually impaired flies. On the last day before the shift, temperature was switched to constant 25°C, and the dark phase was extended by 6 hr. The new regime was kept for 7 days, and subsequently the flies were released into DD for another 5 days. Light pulse (LP) experiments were performed in light- and temperature-controlled incubators as described [10]. Flies were taken out of the incubator during the dark phase (ZT15 or ZT22) on the last day of an LD cycle and transferred to an incubator with lights on. LPs were given for 30 minutes after which the flies were returned to the original incubator. Non-pulsed control flies were kept in the original incubator for the entire experiment.

Plotting of behavioral activity, period calculations, and circular phase plots for the LP experiments were performed using a signal-processing toolbox [53] implemented in MATLAB (MathWorks, Natick, MA, USA). Phase quantification for determining re-synchronization kinetics to shifted LD cycles was done as described in [11] using a custom made Excel macro and the results were plotted in R (R Foundation, Vienna, Austria).

### Immunohistochemistry

Comparison of PER levels in the different neuronal groups and time points was performed as previously described [7]. Briefly, flies were entrained to LD and temperature cycles for four days, flowed by 6 hr delayed LD at constant 25°C as described above for the jetlag experiments. On day two of the phase-delayed LD cycle flies were collected at the indicated time points and fixed in 4% PFA for 2 hr 30 min at room temperature (RT). Once fixed, the flies were washed 1 hr at room temperature with 0.1 M phosphate buffer (pH 7.4), 0.1% Triton X-100 (PBS-T). Brains were dissected in PBS and blocked for 2 hr with 10% goat serum in 0.5% PBS-T. The primary antibodies used were pre-absorbed rabbit anti-PER (1:5000) and monoclonal mouse anti-PDF C7 (DSHB, 1:200). Secondary antibodies were goat anti-rabbit AlexaFluor 488 and goat anti-mouse AlexaFluor 647 (Molecular Probes, 1:500). Brains were mounted in Vectashield (Vectorlabs), and imaged using a Leica SP8 confocal microscope. Quantification of pixel intensity of mean and background staining in each neuronal group was measured using FIJI [54]. For each cell three measurements were taken, as well as 3 measurements of the background of the corresponding slice. Data were analysed and plotted using R. After background subtractions, measurements were normalized to the values of each of the neuronal groups obtained for *norpA*^*P41*^ *rdgA*^*1*^;;*cry^02^* at ZT22. The images shown in Figure 4 C were processed with GIMP.

### Statistics

Data are presented as mean ± S.E.M. Statistical analysis in Figure 4 was performed in R using one-way ANOVA followed by Tukey’s test.

### Sequencing of the *rdgA* coding sequence

In order to cover for possible changes between alleles, both *rdgA A* and *rdgA C* were sequenced from wild type (*w1118*) and *rdgA*^*1*^ mutant flies. cDNA was generated from 20 heads following the protocol from [55]. Briefly, the heads were collected in 2ml RNAlater (Ambion) with 100μl 0.1% PBST to help penetration. mRNA was extracted following the RNEasy kit (QIAGEN). The RNA was eluted in RNase-free water and used for cDNA synthesis following the instructions of the Reverse Transcription Reagents Kit (Applied Biosystems) in 20 μl reactions. PCR with primers specific for both alleles (Table S2) were performed on both the wild type and mutant samples, and the products prepared for sequencing. The obtained sequences were compared to one another and to the published sequences of *rdgA* [32].

### ERG recordings

Electroretinograms (ERG) were recorded as described previously (e.g. [7,56]) from young (1-5 days old) dark-reared flies of either sex immobilised with low melting point wax in truncated pipette tips. Recordings were made with low resistance (~10 MΩ) glass microelectrodes filled with fly Ringer (140 mM NaCl, 5 mM KCl, 1.5 mM CaCl_2_, 4 mM MgCl_2_) inserted into the retina, with a similar electrode inserted into the head capsule near the ocelli as reference. Light stimulation came from a “warm white” power LED (Cairn Research) delivered to within ~ 5 mm of the eye via a liquid-filled light guide (diameter 5 mm). Intensity was controlled with neutral density filters (Comar Optics UK).Maximum intensity of the unattenuated stimulus was equivalent to approximately 10^7^ effectively absorbed photons per second per photoreceptor. All flies were on a white-eyed (*w^1118^*) background.

### Optical neutralization

Rhabomere integrity was assessed in young (1-2 day old) red-eyed flies as previously described (e.g. [24,57]) by observing rhabdomere patterns with antidromic white illumination under an 40× oil immersion objective focused below the cornea of decapitated heads fixed to a microscope slide using clear nail varnish. In wild-type flies the classic pattern of 7 rhabdomeres in a trapezoidal/hexagonal array are clearly visible in every ommatidium; retinal degeneration is apparent from distorted patterns and/or missing rhabdomeres.

### Spectophotometric measurements of rhodopsin

Relative rhodopsin content was estimated as previously described (e.g. [58]) by exploiting the spectral properties of the two, photointerconvertible thermostable states of the fly visual pigment: rhodopsin (R), absorbing maximally at 480 nm and metarhodopsin (M) absorbing maximally at 570 nm. Long wavelength light scattered back out of the eye is more effectively absorbed by M than it is by R. To measure the back-scattered light, intact flies of different genotypes on white-eyed backgrounds were mounted in truncated pipette tips (as for ERGs) and imaged with a 20× air objective on an inverted Nikon TMS microscope. Photoequilibrating blue light (470 nm) was first delivered to generate ~70% M and then yellow/green light (540 nm) was delivered to convert M back to R. The back-scattered light, which was measured with a photomultiplier (Cairn Research UK), increased exponentially as M was photoreconverted to R, and the relative increase provides a measure of the concentration of visual pigment (Fig. S1).

