## Supplemental Data for "Less is more: Light sampling via throttled visual phototransduction robustly synchronizes the *Drosophila* circadian clock in the absence of Cryptochrome"

### Figure S1

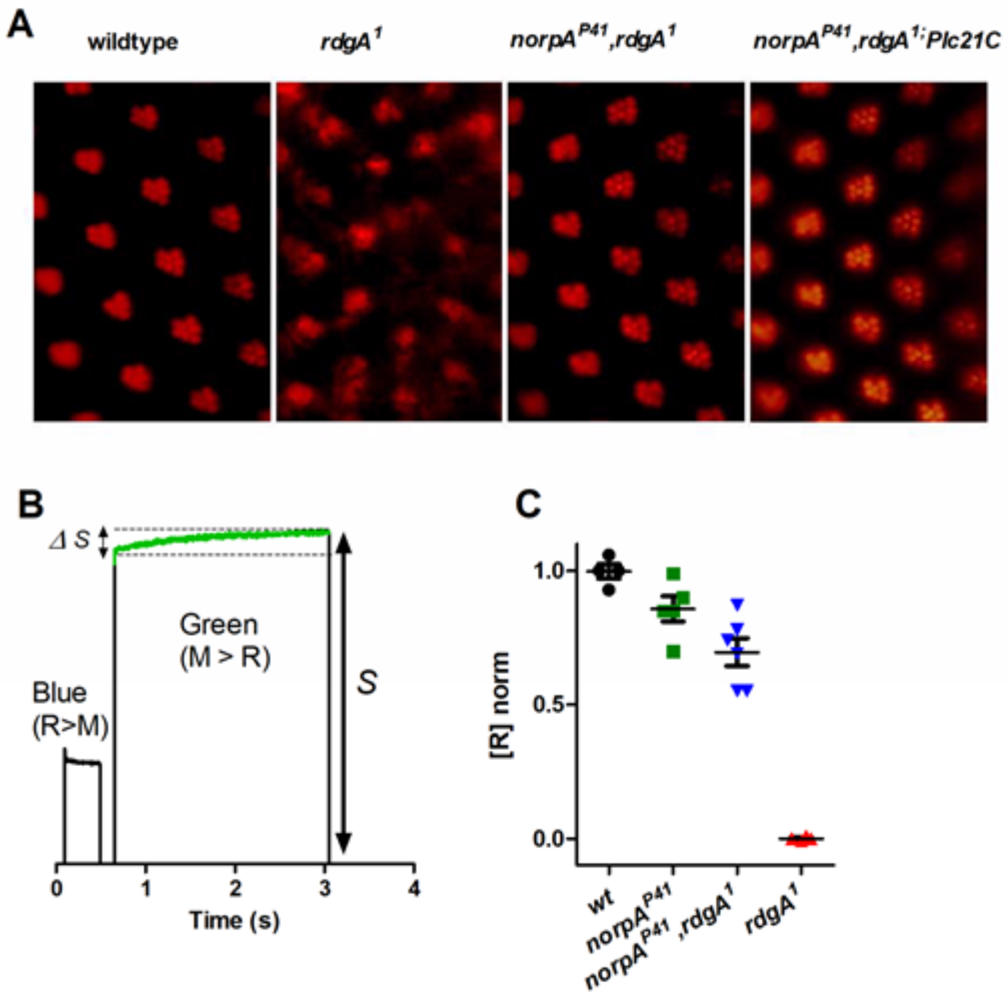

A
wildtype
rdgA1
norpAP41,rdgA1
norpAP41,rdgA1;Plc21C
B
ΔS
Green
(M > R)
Blue
(R>M)
S
0
1
2
3
4
Time (s)
C
1.0
0.5
0.0
[R] norm
wt
norpAP41
norpAP41,rdgA1
rdgA1

**Figure S1. Full rescue of *rdgA$^{1}$*-induced photoreceptor degeneration by *norpA$^{P41}$*.** A) Rhabdomeres visualized by optical neutralization from red-eyed wild-type, *rdgA$^{1}$* single mutant, *norpA$^{P41}$,rdgA$^{1}$* and *norpA$^{P41}$,rdgA$^{1}$;Plc21C$^{P319}$* double and treble mutants. Heads of dark-reared young flies were illuminated with antidromic white light under oil immersion. B) Spectrophotometric method for determination of relative rhodopsin content in vivo in white-eyed flies (Randall et al., 2015): first blue light (470 nm) is delivered to convert the majority of (blue-absorbing) rhodopsin to green/yellow absorbing metarhodopsin. Subsequently, green light (540 nm) is delivered to convert M back to R) and the light scattered back out of the retina is measured (S). As M is photoreisomerised, the green light backscattered out of the retina increases with an exponential time course representing the rate of photoreisomerisation (M>R). The relative increase in back-scatter (ΔS/S) is a measure of the rhodopsin content. C) ΔS/S values normalized to wild-type (*w$^{1118}$*) give relative rhodopsin (R) content. There was no detectable increase in backscatter (i.e no detectable rhodopsin) in *rdgA$^{1}$* mutants, but levels were near normal in *norpA$^{P41}$* and *norpA$^{P41}$,rdgA$^{1}$*.

### Figure S2

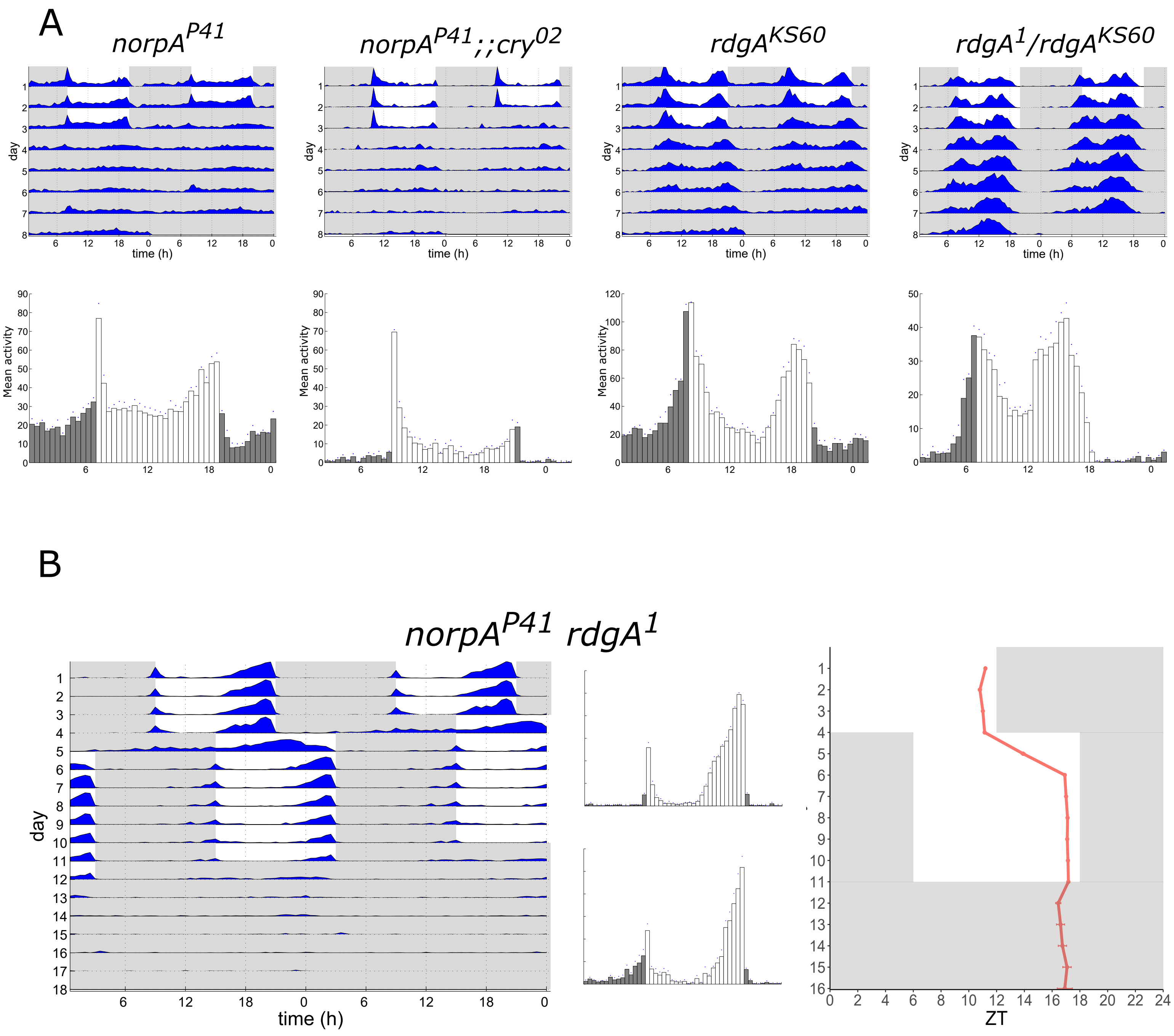
A
norpA^P41
norpA^P41;;cry^02
rdgA^KS60
rdgA^1/rdgA^KS60
day
time (h)
Mean activity
B
norpA^P41 rdgA^1
day
time (h)
ZT

**Figure S2. Control genotypes for resynchronization experiments to 6-hr delayed LD cycles.** (A) Representative actograms (top) and histograms (bottom) of 16 flies of the indicated genotypes subjected to 3 days in LD at 25 °C followed by 5 days in DD at 25°C. (B) Results for *norpA*$^{P41}$ *rdgA*$^{1}$ double mutants subjected to similar "jetlag" conditions as in Figure 3, where flies were exposed to 4 days of combined LDTC, followed by a 6 hr delay of the LD cycle at constant 25°C. Average actogram (left), histograms (middle) and phase quantification (right). The histograms represent the first 3 days before the phase shift (top) and the last 3 days after the shift (bottom).

### Figure S3

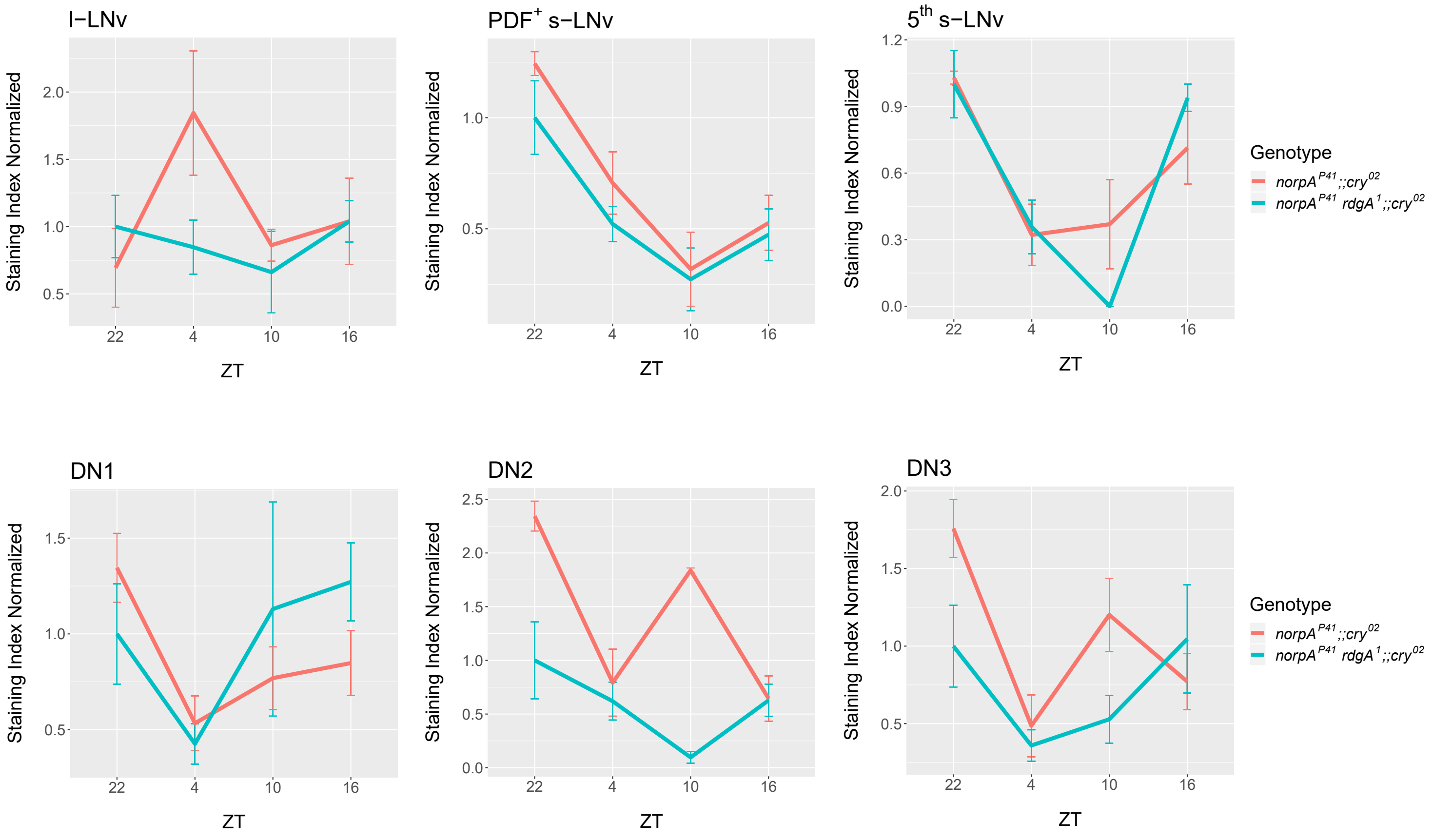
l−LNv
PDF+ s−LNv
5th s−LNv
DN1
DN2
DN3
Staining Index Normalized
ZT
22
4
10
16
Genotype
norpA P41;;cry02
norpA P41 rdgA 1;;cry02

**Figure S3**. **Improved PER synchronization in clock neurons of *norpA$^{P41}$rdgA$^{1}$ cry$^{02}$* treble, compared to *norpA$^{P41}$ cry$^{02}$* double-mutants.** Quantification of PER expression in individual clock neuronal groups (A), and in all groups combined (B) for the indicated genotypes. Brains were dissected at the indicated time points on day 2 of a 6-hr delayed LD cycle in a jetlag experiment (e.g., see Figure 3A). Between 3 and 6 brain hemispheres were analyzed for each genotype and time point.

### Figure S4

A

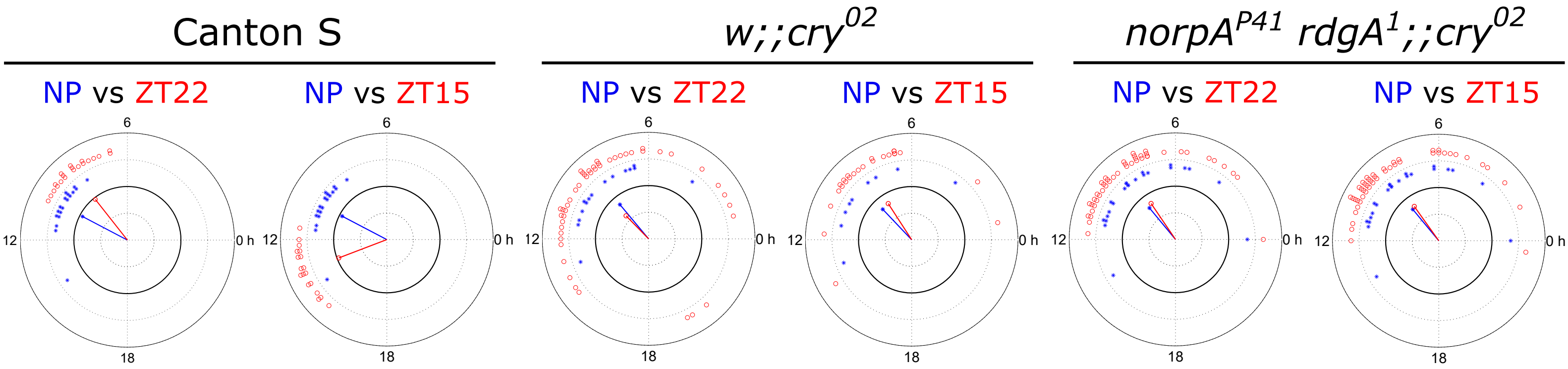
Canton S
w;;cry02
norpAP41 rdgA1;;cry02
NP vs ZT22
NP vs ZT15
NP vs ZT22
NP vs ZT15
NP vs ZT22
NP vs ZT15
6
12
0 h
18

B

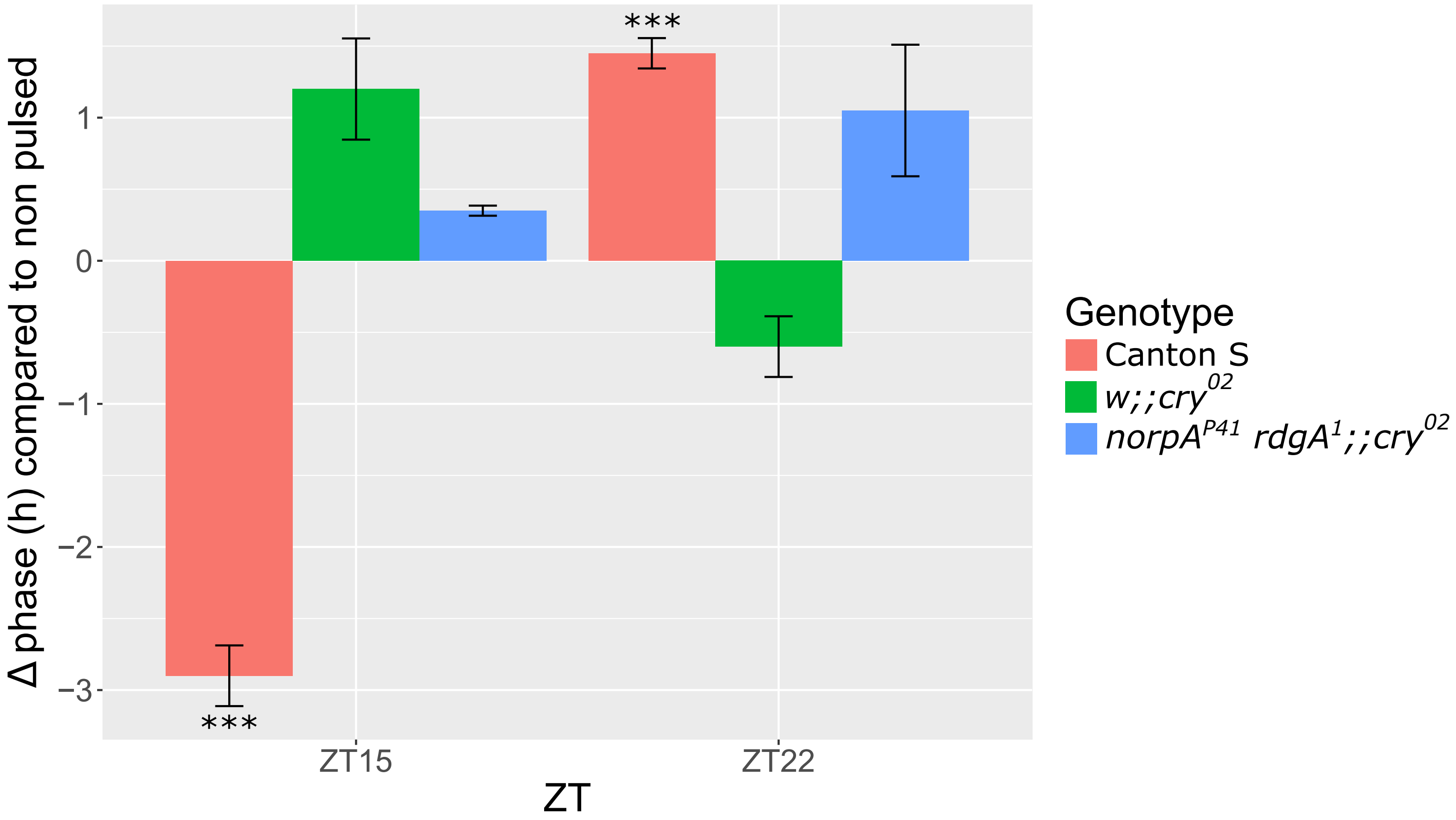
Δ phase (h) compared to non pulsed
1
0
−1
−2
−3
***
***
ZT15
ZT22
ZT
Genotype
Canton S
w;;cry02
norpAP41 rdgA1;;cry02

**Figure S4. Visual system function in *norpA*$^{P41}$ *rdgA*$^{1}$ double mutants is not sufficient for behavioral clock synchronization to 30 min Light pulses.** Comparison of peak activity phase after a 30 min light pulse (LP) given at ZT15 and ZT22 in subsequent constant conditions. (A) Circular phase plots showing the phase of non-pulsed (blue) and pulsed flies (red) of the indicated genotypes at ZT22 (left) and ZT15 (right) 1 day after the pulse. Dots represent phase of individual flies. The direction of the vector indicates the mean phase of a genotype and the magnitude of the vector indicates the coherence of the group (variance around the mean) (Levine et al., 2002). (B) Bar graph showing the difference of peak activity of the indicated genotypes compared to that of non-pulsed flies of the same genotype. Error bars represent SEM. Δ phase and statistical significance was determined using circular statistics. Canton S, non-pulsed n=34, ZT22 n=40, ZT15 n=41; *w;;cry*$^{02}$, non-pulsed n=37, ZT22 n=69, ZT15 n=68; *norpA*$^{P41}$,*rdgA*$^{1}$;;*cry*$^{02}$, non-pulsed n=50, ZT22 n=67, ZT15 n=67

### Figure S5

| | | | |
|---|---|---|---|
| *D. melanogaster* | 896 | VLPLGTGND | 904 |
| *D. suzukii* | 372 | VLPLGTGND | 380 |
| *D. simulans* | 898 | VLPLGTGND | 906 |
| *Danio rerio* | 394 | VLPLGTGND | 402 |
| *M. musculus* | 453 | VLPLGTGND | 461 |
| *rdgA*$^{1}$ mutants | 896 | VLPLSTGND | 904 |

**Figure S5.** ***rdgA*$^1$ encodes a Glycine to Serine change within the catalytic domain of DAG-kinase.** Comparison of the relevant highly conserved amino acid sequence within the catalytic domain of *rdgA* encoded DAG-kinase between different *Drosophila* species, and the orthologs in zebrafish (*Danio rerio*) and mouse (*Mus musculus*) and the mutant *Drosophila melanogaster* DAG-kinase encoded by *rdgA*$^1$. Numbers represent the position in the protein sequence according to NCBI.

**Table S1. Free running behaviour of the genetic variants used in this study.** RS: rhythmic strength. All values represent the average ± S.E.M.

| Genotype | % Rhythmicity | Period | RS | n |
|---|---|---|---|---|
| Canton S | 97.8 ± 1.8 | 24.3 ±0.1 | 3.3 ± 0.3 | 38 |
| *norpA*$^{P41}$ | 70.3 ± 3.2 | 23.4 ± 0.2 | 2.8 ± 0.2 | 47 |
| *w;;cry*$^{02}$ | 83.3 ± 2.4 | 23.6 ± 0.4 | 2.0 ± 0.2 | 30 |
| *norpA*$^{P41}$*;;cry*$^{02}$ | 64.2 ± 3.6 | 24.1 ± 0.3 | 2.5 ± 0.4 | 50 |
| *rdgA*$^{1}$ | 85.1 ± 2.7 | 23.6 ± 0.2 | 3.2 ± 0.3 | 90 |
| *rdgA*$^{KS60}$ | 76.8 ± 1.4 | 24.4 ± 0.2 | 2.4 ± 0.2 | 43 |
| *rdgA*$^{1}$*/ rdgA*$^{KS60}$ | 100 | 23.5 ± 0.1 | 4.7 ± 0.1 | 31 |
| *rdgA*$^{1}$*;;cry*$^{02}$ | 69.4 ± 1.4 | 23.6 ± 0.6 | 2.5 ± 0.3 | 90 |
| *norpA*$^{P41}$ *rdgA*$^{1}$ | 76.7 ± 1.6 | 23.8 ± 0.2 | 3.1 ± 0.4 | 44 |
| *norpA*$^{P41}$ *rdgA*$^{1}$*;;cry*$^{02}$ | 93.7 | 23.9 ± 0.2 | 2.5 ± 0.3 | 32 |
| *norpA*$^{P41}$ *rdgA*$^{1}$ *;PLC21C*$^{P319}$*;cry*$^{02}$ | 9.6 ± 1.1 | 23.2 ± 0.3 | 1.8 ± 0.1 | 34 |
| *norpA*$^{P41}$ *rdgA*$^{1}$ *;Gq*$^{V303D}$*;cry*$^{02}$ | 28.6 | 23.6 ± 0.4 | 1.8 ± 0.6 | 8 |

**Table S2. List of primers used for sequencing of the *rdgA* CDS**

| | |
|---|---|
| Primers to sequence the common regions between alleles | |
| rdgA primer 1 F | AAGCAGTGCGGCAAGTTCTTT |
| rdgA primer 1 R | ACAATGACCGGGATTACCTCC |
| rdgA primer 2 F | GCCCGATTGGACGGAGAATG |
| rdgA primer 2 R | CCGCCTGCTTCATAGGAAAGAAC |
| rdgA primer 3 F | AGCACTATTGGAAGCCCACC |
| rdgA primer 3 R | TTTGCACCGCCTGCTTCATA |
| rdgA primer 4 F | TTCCTATGAAGCAGGCGGTG |
| rdgA primer 4 R | CGATCGACATGGTCATCGGT |
| rdgA primer 5 F | CTCTCGCAATGGGTGACGC |
| rdgA primer 5 R | ACTGCTCGTACTGTTGCATGG |
| rdgA primer 6 F | TAAACATTCCCAGCTATGGCG |
| rdgA primer 6 R | AGCACCAATCCTGTGAGACC |
| rdgA primer 7 F | AAGGCACTGATGCTGTCGAA |
| rdgA primer 7 R | GCTCCAGTATAGGGCTCTTCC |
| rdgA primer 8 F | GAGTGCATTAAAAGCGGCCA |
| rdgA primer 8 R | TTTGCACCGCCTGCTTCATA |
| Specific primers for allele A | |
| rdgAA F1 | TCGAATGCCAAAACGGAGGA |
| rdgAA R1 | TAGGAAAGAACTTGCCGCACT |
| rdgAA F2 | TGTGGCATAACCTTCACGCT |
| rdgAA R2 | GGTACGCCCGTCTGCATAG |
| rdgAA F3 | GACGAGGCGCGTTGTTCTA |
| rdgAA R3 | TGTTGGACACCAGAAGCCAG |
| rdgAA F4 | CCAGAGAGCCGACCAAGAAC |
| rdgAA R4 | AGGCCGATGATGAGGATTGC |
| rdgAA F5 | GATCACGCACTCCAACAGGT |
| rdgAA R5 | TGCTGTAGCTCAAAGCCGTT |
| rdgAA F6 | ACGGCTTTGAGCTACAGCAA |
| rdgAA R6 | CTCGGACTGGCAAACCCATA |
| Specific primers for allele C | |
| rdgAC F1 | GCAGTGATACAATCTCTGGGC |
| rdgAC R1 | ACGTTTCCCTCATCGTCCAC |
| rdgAC F2 | AATTGTGCAAGTGGCGTTCG |
| rdgAC R2 | CGTGTCTGCGGCTGTCATTA |
| rdgAC F3 | GATTGCTGCACGCTGTTCG |
| rdgAC R3 | GGCCGCTTTTAATGCACTCC |
| Primers for genomic confirmation of the mutation | |
| rdgA1 F | CGAGCGTTCATTGTGAAGCC |
| rdgA1 R | TGGGGAGGGGTGAGTAATCTT |
